# Torsion angles to map and visualize the conformational space of a protein

**DOI:** 10.1101/2022.08.04.502807

**Authors:** Helen Mary Ginn

## Abstract

Present understanding of protein structure dynamics trails behind that of static structures. A torsion-angle based approach, called representation of protein entities (RoPE), derives an interpretable conformational space which correlates with data collection temperature, resolution and reaction coordinate. For more complex systems, atomic coordinates fail to separate functional conformational states, which are still preserved by torsion angle-derived space. This indicates that torsion angles are often a more sensitive and biologically relevant descriptor for protein conformational dynamics than atomic coordinates.

## 2 Introduction

Researchers can now deliver structure solutions for protein targets at remarkable speed, evidenced by the exponentially growing number of Protein Data Bank (PDB) [1] depositions. This database has reached sufficient size to support the development of AlphaFold [2] to predict protein structure, which is distinct from, but may re-open efforts towards protein folding prediction [3]. This progress is tempered by the smaller strides made towards understanding allosteric regulation and subtle conformational shifts, which often govern enzyme regulation, activity and signalling [4]. For example, we still do not know whether cryo-cooling samples for data collection, which underpins the most experimental structure determinations, has a deleterious effect on biologically relevant information [5–8], and whether the quest for an interpretable high-resolution map is worth the trade-off in trapping a single conformational state potentially less relevant to biological function. In short, our understanding of dynamics, and hence the essence of biological activity, lags far behind our understanding of static structures.

Beyond experimental data collection, we have computational tools to analyse structures. Individual crystal structures can be analysed using tools such as ECHT [9], which hierarchically removes TLS-derived modes of motion from incrementally smaller atom groups expected to harbour collective motion, from whole-polypeptide chains to domains, secondary structure and individual residues. Al-ternatively, qFit3 [10], which fits multiple conformations to electron density by sampling variations of torsion angles, can sense and explain dynamic motion in the structure. Crystal structures comprise multiple different states of the protein, and xtrapol8 allows for automatic estimation of the proportion and subtraction of a known state of the protein from the electron density [11]. Binding events of small molecules can be detected from multi-datasets using PanDDA which is regularly used in drug- and fragment-screens [12]. 2D and 3D classification in cryo-electron microscopy is also able to select enriched conformations from a heterogenous set of particles [13]. Ensemble refinement combines molecular dynamics (MD) and fitting to experimental data [14]. MD has also been adapted to crystal lattices and supercells, as previously reviewed [15]. Tools such as normal mode analysis [16] and elastic network models [17] have a lower computational cost than MD. Diffuse scattering is a potential experimental source of dynamic information, but application to biomolecules is exceptionally difficult [18]. Despite these tools, we continue to struggle to accurately describe and track subtle conformational states in proteins beyond the easily recognisable large domain or secondary structure movements and rearrangements. These difficult-to-visualise subtle shifts are nevertheless vital for, and intrinsically bound to, activity, efficacy and regulation. In this paper, I introduce a method capable of discerning subtle conformational changes across the entirety of a biomacromolecule. This approach uses torsion angles to describe conformational states, a more biologically relevant quantity than atomic coordinates: although (x, *y, z*) coordinates are the simplest method to define an atom’s position, atomic motion does not naturally align with this parameter space. Proteins naturally sample the conformational space by variation in torsion angles, which have a significantly lower energy barrier than changing bond lengths and angles. This powerful space shows correlations with resolution of the dataset, data collection temperature, space group and bound ligand, that goes beyond what is achievable using atomic coordinates.

This multi-dataset analysis is named Representation of Protein Entities (RoPE). The product of this analysis is a representation of protein conformational space (hereby referred to as RoPE space). What were previously considered subtle shifts in the proteins conformational state are starkly separated in RoPE space and provide a means for classification and comparison. RoPE is a potentially powerful tool that can also be used to drive experimental decision-making to help experimentalists obtain data suitable to address complex biological questions dominated by protein dynamics. The algorithm is contained within a software package and available as an open source application (https://rope.hginn.co.uk).

## 3 Results

### Calculation of RoPE space

PDB structures from PDB-redo [19] were loaded and structures recalculated using ideal bond length and angle geometry, and torsion angle estimates from each set of four connected atoms. This produces a nonsensical polypeptide path (Figure 1A) due to the propagation of many small bond length and angle errors. Vagabond refinement [20] was used to obtain optimal estimates of torsion angles, by restoring the polypeptide path to that of the original structure (Figure 1B). Sequences were extracted from the PDB and aligned in order to identify matching torsion angles across all PDB files. All torsion angles, excluding those for hydrogen atoms, contribute to the calculation of the RoPE space, as described in the Methods. Singular value decomposition (SVD) was carried out on the torsion angle difference vectors. Figure plots were generated by manual rotation of the first three axes to reveal the pertinent information and then projected into 2D. Therefore, the X and Y axes are some combination of the first three principle components, best showing the variation with respect to the metadata. Each data point represents one polypeptide chain. This workflow is summarised in Figure 2.

**Figure 1:**
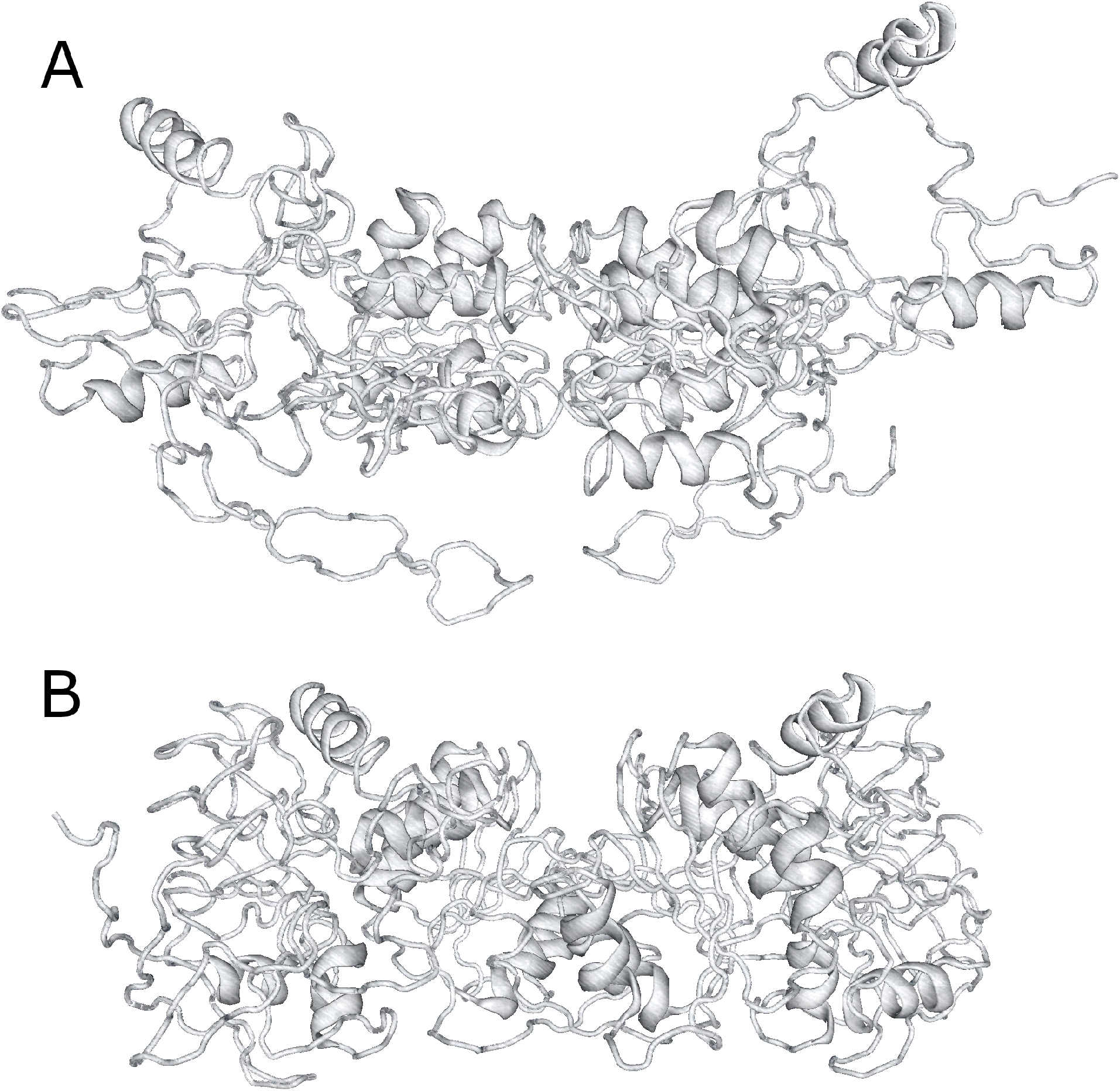
(A) alcohol dehydrogenase using torsion angles directly calculated from sequential sets of four atoms and expected bond length and angles as defined in the literature. (B) after optimisation of torsion angles, alcohol dehydrogenase follows the path of the original structure.

**Figure 2:**
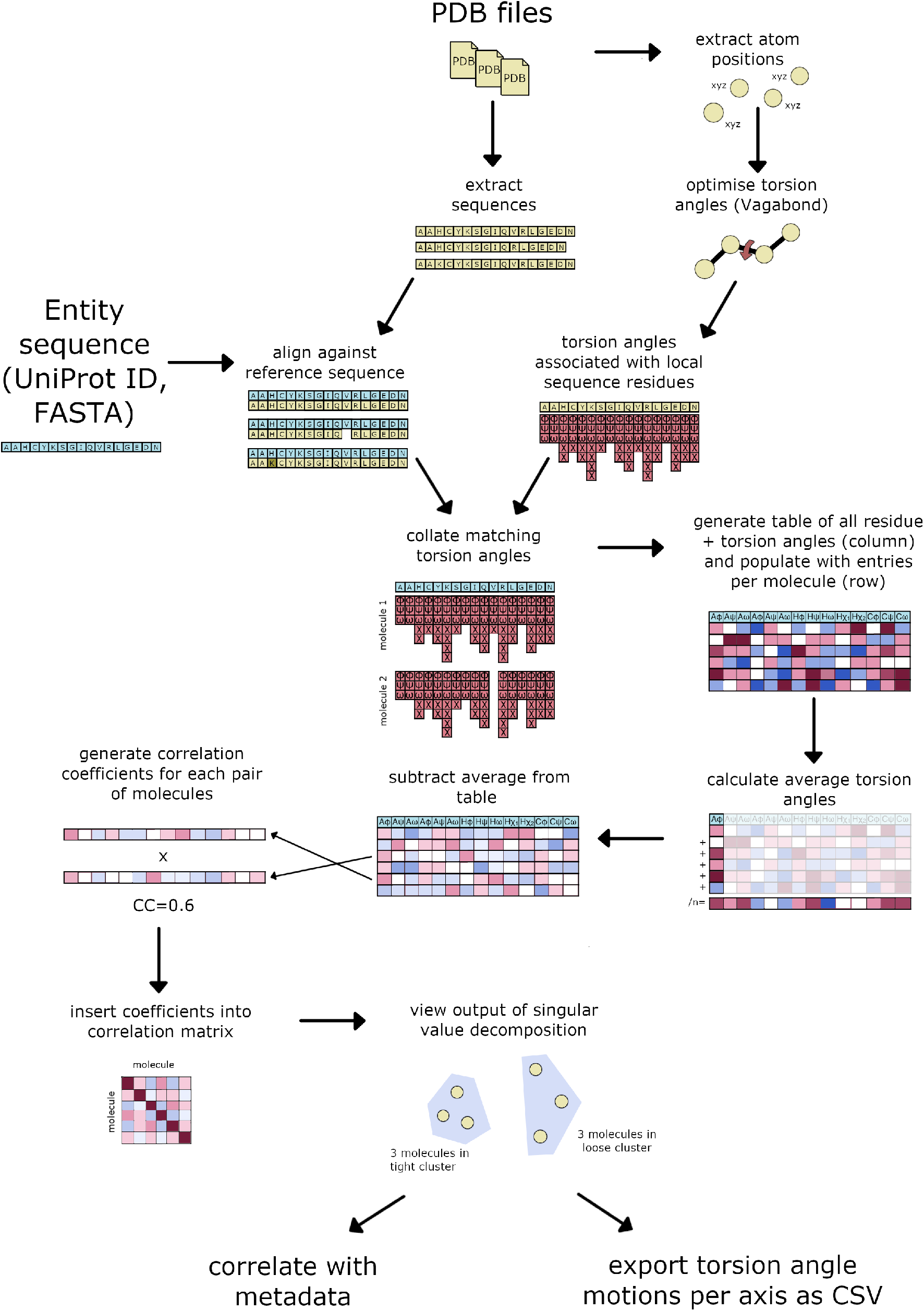
Workflow showing how input data is handled in RoPE. PDBs are used to extract both atom coordinates and sequences. A reference sequence is supplied externally, to which each PDB sequence is aligned. Atom coordinates form the basis of torsion angle refinement. Sequences and calculated torsion angles are combined to facilitate clustering.

In order to provide a comparison, conformational spaces were also calculated using atomic coordinates, after alignment to a reference structure, as explained in Methods. The choice of reference structure is a confounding factor, as the quality of the alignment strongly affects the results. Such spaces below are calculated for comparison, for which a suitable (but not necessarily optimal) reference protein has been chosen. This decision is entirely avoided by using torsion angles, which report only on the internal relationships in the protein and does not require a reference. The following cases are reported in order of increasing complexity, where torsion angles remain useful as atomic coordinates become increasingly incapable of separating conformational states.

### Comparison with atomic coordinates

A time-resolved experiment on the dynamics of photoexcitation of carboxymyoglobin to initiate loss of carbon monoxide consists of timepoints between −0.1 ps and 150 ps from the point of photoexcitation [21], with a time resolution on the scale of femtoseconds. The experiment produced very small shifts in the backbone of the myoglobin, with a reported maximum RMSD of 0.11 Å between the first and last timepoint [21]. A correlation matrix showing similarity of torsion angle vectors (Figure 3A) shows the correlation of each structure is higher with its temporal neighbours up to 3 ps. The last three timepoints, 10 ps, 50 ps and 150 ps, are beyond the biologically relevant timescales for CO dissociation in myoglobin, and neither did these strongly correlate with any other timepoints. This correlation matrix corresponds to the SVD space as shown in Figure 3B. For these small motions, linear changes in torsion angles almost totally correspond to linear atomic coordinate motions. Although this correspondence will break down for larger motions through violation of the small angle approximation, atomic coordinate analysis produces a similar conformational space in the case of myoglobin (Figure 3C). In this case, one may ask: what is the benefit of using torsion angle space over atomic coordinates? One advantage is that the superimposition of structures is not simply required, and therefore choice of reference structure is avoided. Another advantage of using torsion angles, even for small motions, is that each principle axis can be mapped onto the torsion angles in the structure, in order to highlight the hinges in motion during the trajectory of the protein (Figure 3D-E).

**Figure 3:**
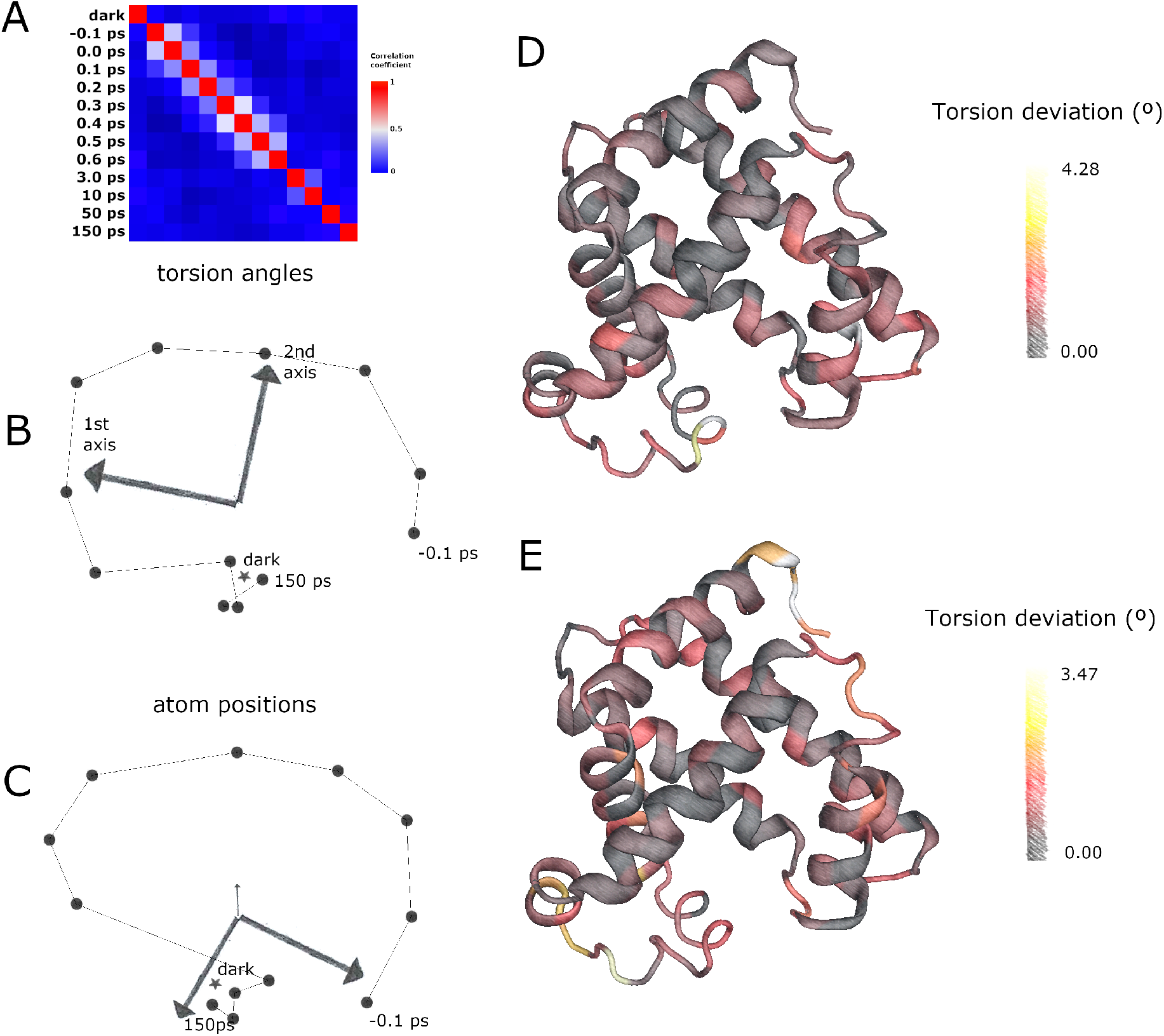
(A) correlation matrix between torsion angle vectors for each myoglobin dataset. (B) corresponding RoPE space for myoglobin dataset. Dark structure marked as a star, −0.1 ps to 150 ps timepoints marked as a line series. 1st and 2nd principle axes are marked. (C) conformational space generated using atomic coordinates. (D-E) cartoon view of myoglobin with colour of each residue proportional to the sum of deviation of its Ramachandran angles, for torsion angles corresponding to the first and second principle axis respectively.

### Variation with temperature

Variation of temperature during data collection can also produce atomic motions and torsion angle motions that allow separation between room temperature and cryocooled structures. From 459 crystal structures, 515 molecules of bovine trypsin were extracted from 459 crystal structures, with some containing multiple copies in the asymmetric unit. Torsion angles were optimised for all structures collected between 1982 and 2022 which had a corresponding entry in the PDB-redo database [19]. Recognition of cryo- and RT by overlaying structures by eye is prohibitively difficult. Conformational spaces were either calculated from torsion angles to produce a RoPE space (Figure 4A) or from atomic coordinates (Figure 4B), which show a strong degree of equivalency. In both cases there is a largely clean separation of room temperature and cryo-cooled structures, with a slightly improved separation when using torsion angles to generate a RoPE space.

**Figure 4:**
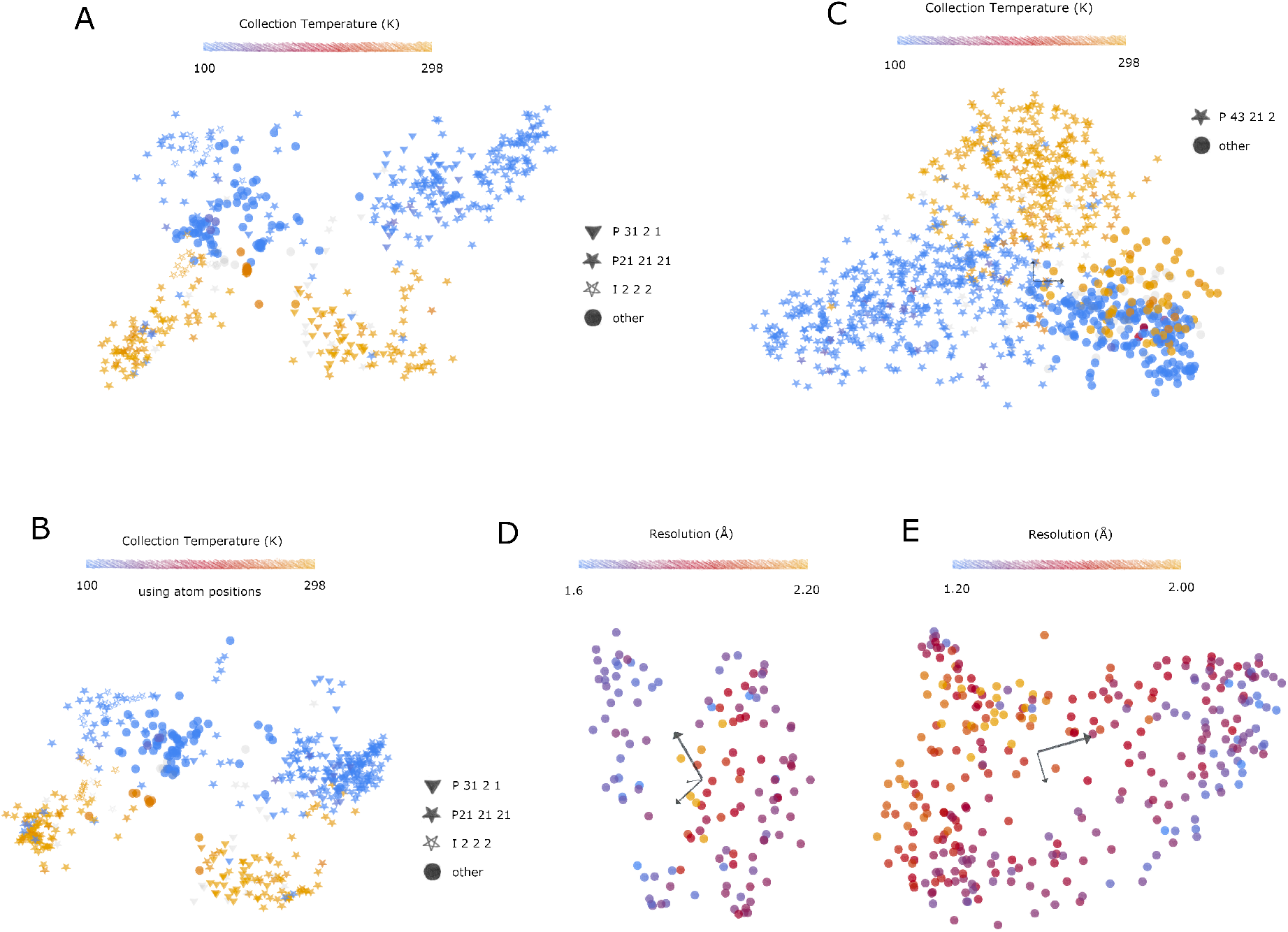
RoPE spaces for various proteins where each point represents a molecule from a PDB file. (A) trypsin (Uniprot ID P00760, 459 structures, 511 molecules) showing split on collection temperature. (B) conformational space generated using atomic coordinate differences for trypsin. (C) lysozyme (Uniprot ID P00698, 939 structures, 1038 molecules) with the majority space group P43212 marked as triangles. (D) BAZ2B, drug fragment screen for PanDDA analysis, 132 structures/molecules, coloured by resolution. (E) DNA cross-link repair protein 1A, deposition ID G_1002036, 312 structures/molecules, coloured by resolution. (A-C) light grey points indicate that the corresponding metadata were not available, as the Protein Data Bank requested less information during deposition for earlier PDBs.

Notably in Figure 4A, there were eight outlier structures where reportedly cryo-cooled molecules were present in the room temperature cluster. Six of these cases were contradicted by the published methods, stating that they were indeed collected at room temperature [22, 23], suggesting that the wrong metadata were deposited in the PDB. Other publications did not specify the collection temperature directly [24,25]. However, another paper confirmed data collection for two other trypsin outliers was carried out at 100 K [26]. This suggests that these outliers were largely caused by erroneous annotation in the PDB, and that these conformational spaces are therefore powerful enough to identify these metadata errors, and the collection temperature could be accurately estimated from a set of atomic coordinates even without the need to factor in the space group or unit cell.

Not all structures have a similar RT/cryo-cooled distinction in RoPE space, and for most there is insufficient sampling to draw a conclusion. Some splits are less extreme; the RT and cryo-cooled regions of lysozyme overlap more than for trypsin (Figure 4C). In both cases however the torsion angle difference associated with the temperature change is essentially preserved across space groups. RoPE space can therefore be used to determine on a per-protein basis if the biological interpretation of the structures is likely to be perturbed by the RT/cryo-cooled conformational split.

### Variation with resolution

Drug or fragment screens provide an excellent supply of multi-datasets from crystals selected specifically for their homogeneity. Despite this, often large variations in resolution between crystals remain, which are largely unexplained. The RoPE space for a two drug or fragment screens was determined. BAZ2B (Fig. 4D) [12] and the DNA crosslink repair protein 1A (Fig. 4E) show a clear dependence of resolution with a molecule’s position within the RoPE space. In each case, there is a tendency for the high resolution structures to coalesce some of the edges of the occupied space. This suggests that molecules that lie in the middle of the RoPE space may actually be sampling a wider distribution of neighbouring conformations in the crystal, blurring the electron density and decreasing the reported resolution. An alternative explanation, however, is that high-resolution structures are intrinsically better defined. Correlation coefficients are therefore naturally higher and more accurate, and will produce horseshoe effects [27] when combined with poor estimations of correlation with lower resolution structures. Evidence against this is demonstrated by atomic coordinate-derived conformational space and torsion angle-derived RoPE spaces of haem subunits. In this case, atomic coordinates produce a conformational space of the alpha (Figure 5A) and beta (Figure 5B) subunits with a poor correlation with resolution and are in fact subject to extreme horseshoe effects regardless of resolution. Conversely, the RoPE spaces of the alpha (Figure 5C) and beta (Figure 5D) subunits do not suffer from horseshoe effects. There is nevertheless a far stronger dependency of resolution on its neighbours in torsion angle space. This shows that resolution is best interpreted in torsion angle space and not atomic coordinates. The left and right lobes correspond to an adjustment of the haem group relative to its protein scaffold which is consistent with the expected change between the deoxy- and oxyhaemoglobin states [28].

**Figure 5:**
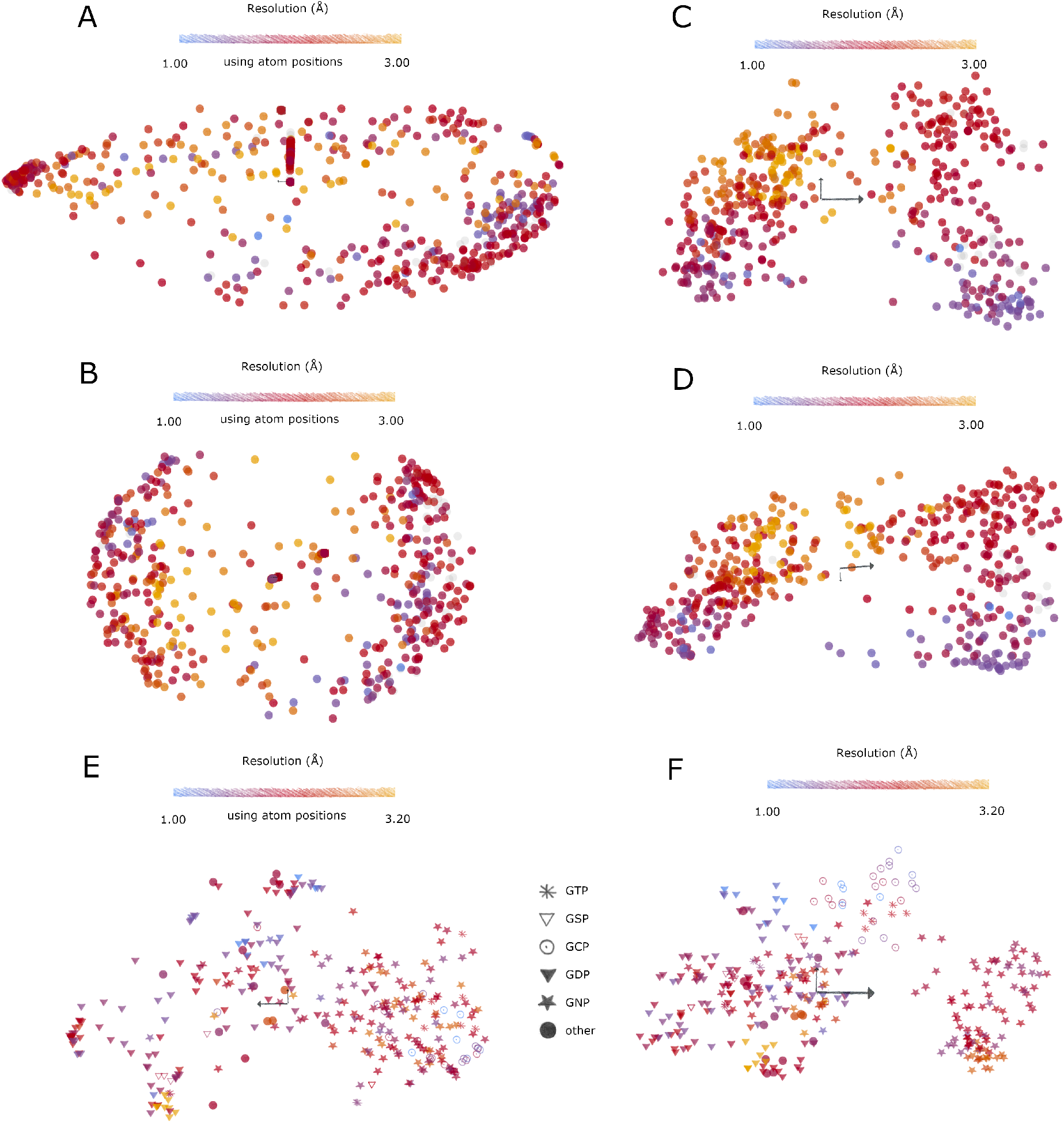
Atomic coordinate-derived conformational space for the (A) alpha subunit of haem (Uniprot ID P69905, 498 molecules) and (B) the beta subunit of haem (Uniprot ID P68871, 497 molecules). Corresponding RoPE spaces using torsion angles for (C) the alpha subunit and (D) the beta subunit, respectively. (E) atomic coordinate-derived conformational space for kRas, and (F) RoPE space using torsion angles for kRas.

### Variation with bound ligand

Protein conformation also responds to the identity of ligands bound. One such example is kRas, of the Ras superfamily of proteins, which binds guanine nucleotides. kRas in the inactive state, which binds GDP, are stable enough to crystallise. Active forms of kRas auto-hydrolyse the ligand, but can be stabilised through inactivating mutations or using non-hydrolysable analogues of GTP, namely GSP, GNP, and GCP, for which the terminal phosphate is bridged by a sulphur, nitrogen or carbon respectively. This allows trapping of the active states of kRas, which are known to vary as a function of bound ligand both in occupancy and kinetics [29]. The atomic coordinate space shows mediocre separation of the ligand-bound structures. RoPE space (Figure 5E) showed significantly tighter clusters and use of torsion angles is able to distinguish GCP binding from other binding modes (Figure 5F). The separation of this cluster in the atomic coordinate-derived space is not recovered by exploring additional axes of the SVD subspace.

### 4 Discussion

Torsion angles are significantly more sensitive for describing and evaluating protein conformations than atomic coordinate displacement. Linear interpolation in torsion angle space is also straightforward, and leads to structures which do not violate geometric expectations, whereas linear interpolation in atomic coordinates will rapidly deviate from expected bond length and angles. This makes it a more comfortable parameter space to use for expressing protein motion, which will be exploited in future studies. This paper demonstrates that although atomic coordinates can be used to derive a useful conformational space, it fails in many cases to scrutinise conformations. Torsion angles remain a robust clustering device and can distinguish states which atomic coordinates cannot. It is not unreasonable to assume allosteric mechanisms and conformational states in the catalytic cycle of metabolic enzymes will be better-suited to analysis in RoPE space. Future analyses will work towards establishing why proteins are confined to their particular RoPE spaces, each of which have their own idiosyncrasies, and how proteins may find a path in torsion angle space from one conformational state to another.

For trypsin, one of the largest divides is on data collection temperature, and points to the nature of the potential artefact caused by cryo-cooling crystals. RoPE space may provide a computational assay for determining whether cryo-cooled crystals are acceptable alternatives to using room temperature crystals on a case-by-case basis. However, the preservation of other linear independent motions may prove a tolerable trade-off for improved crystal endurance in face of radiation damage, which is a major concern for most protein crystals. This analysis may be a method to establish a reasonable estimation of how cryo-cooling introduces artefacts for a given protein system.

RoPE analysis is not fundamentally limited to X-ray diffraction studies. The analyses in this paper have benefited from standardisation of refinement of X-ray structures by PDB-redo. RoPE is however generally applicable to atomic models regardless of data source. Researchers who trust their atomic coordinates can proceed with data from any experimental method.

### 5 Practial use of the software

In order to demonstrate the utility of the software, a practical example is illustrated using the publicly available datasets of horse liver alcohol dehydrogenase. This uses a multi-dataset of 72 X-ray crystallographic models from the PDB at the time of writing, where a PDB-redo model is available. This discussion largely avoids biological interpretation of the motions in this protein to favour a tour of the important features of using the software, but a review on alcohol dehydrogenase conformations during catalysis [30] is a good accompanying text. This half of the article introduces the most important features of the RoPE software, by instruction of how to explore conformational changes within the subset of alcohol dehydrogenase structures that crystallise in P1.

RoPE is available as a web application at https://rope.hginn.co.uk or downloadable as a Mac OS X executable from the same website. Native applications are preferable. Firefox is the recommended browser to run RoPE, and users should be aware that computation occurs through the browser on the local machine. The code is released under the GPL3 license [31].

RoPE has its own environment, consisting of files, models, entities and metadata. Files are exposed to the system through the project folder on launch of the native, or through the **Menu** > **Load** option in the web application.

The RoPE concept revolves around protein entities, which are defined by a primary sequence. In the case of alcohol dehydrogenase, this is the sequence defined in the UniProt database. From **Protein entities** > **Add entity** > **From UniProt ID**, type in the UniProt ID p00327. This will fetch the sequence. The entity must be named by clicking **Enter…** next to Entity name and then entering an identifier (“**alcohol-dehydrogenase**”), followed by **Create**.

In order to automatically fetch proteins from the PDB, enter the entity by clicking on **alcohol-dehydrogenase** and click **Search PDB** > **Run**. Click **Download** to fetch the PDB files from the PDB-redo server into the local file system, which happens in the background.

Files loaded into the system do not automatically translate into the RoPE definition of models. Models are linked to PDB files and can carry additional metadata. In order to make models, click the **Back** button to return to the **Protein Entities** menu. Click the toolkit symbol in the top right and click **automodel**. This will automatically turn all PDB files within the system into models. It will also automatically associate existing entities with parts of a model if they have over 80% sequence identity.

This is a good time to click back until returned to the main menu, and then click **Menu** > **Save**. The web application will issue an invitation to save a rope.json file to the local filesystem, whereas the native application will save rope.json to the project folder. Note that for generating figures, navigating to **Menu** > **Options** brings up the option of changing the background to pure white.

Once modelling is complete, now the Vagabond algorithm can be run to optimise the torsion angles to best match the atom positions. In order to do this, click on **alcohol-dehydrogenase** and click on **RoPE space**. At the moment all models are unrefined and you will be asked if you wish to refine them. Click **Yes**. This is the right point to schedule a tea break.

Once all proteins have been refined, click the **Back** button to reveal the RoPE space. This is also a good point to save your refinement progress. Each icon represents one monomer of alcohol dehydrogenase. This RoPE space can be rotated with a left mouse-button drag, zoomed with a right-click drag, and panned with a Ctrl+left mouse button drag. This RoPE space currently lacks annotation but it is clear that there is some substructure to the occupied set of torsion angles in alcohol dehydrogenase. Molecules which are close together have similar conformations. We need to know more about what each structure contains, so click **Back** again to return to the entity menu.

Click on **Add to metadata** and then choose from the **Protein Data Bank** in order to automatically download supporting information such as resolution, collection temperature and space group. Note that this uses the model name as a search term, and can fetch information when this matches the PDB code. Once this has downloaded, click on **Add to database** to add this to the known metadata.

Click the **back** button and return to **RoPE space**. It is now possible to add annotations to this space with the additional metadata. The coenzyme-bound, ligand-free holoenzyme of horse liver alcohol dehydrogenase often crystallises in P1, so this can be assigned a different icon. Click on the **rules** menu and click on **(+) new rule**. The options at the top specify what kind of rule to add. To select a discrete value, click on **Change icon**. In order to change what metadata title this responds to, click on **Choose…** and then click **Space group**. Enter the **Options** menu to change the properties of the metadata selection and the icon type. The default is to change the icon type on the basis of having any kind of assignment. However, all our models have a space group of some form, and we only want to select on P1. In order to narrow down the value, click Choose to see the options. On the right hand side is the number of models for each assignment. Choose **P 1** and this will return you to the options menu. Choose an icon type (except for **filled circle**, which is the default) and press **OK**, then **Create**, and then **OK** to see the effect on the RoPE space.

The RoPE space now shows that two of the variations of direction are covered by P1-only space groups which form a V shape. Mouse-over the icons reveals the model name and chain number of each monomer. Each P1 model comprises a dimer, and each direction contains one monomer from each model. Inspection reveals that each monomer of a dimer are in roughly similar positions up each branch of the V. Those at the top of the V have the greatest variation in conformation between monomers (e.g. PDB chains **7uee_A** and **7uee_B** are at the top of the V). On the other hand, P1-structures with minimal separation are found at the pit of the V such as **1qv6_A** and **1qv6_B**.

In order to focus on the P1-structures only, click on the icon legend **Space group = P1** on the right hand side and click **choose group**. This will re-run the RoPE algorithm on the specified subset of structures.

Each direction in RoPE space corresponds to a variation in all torsion angles across the entire molecule. The exact nature of these torsion angle motions is determined by principle component analysis during clustering. This chooses torsion angle motions which show evidence of independent motion between structures, which is naturally limited by the sampling of the conformational space provided by the structures. In order to establish what these torsion angle motions mean, it is possible to draw the motion of a torsion angle on a reference structure. As an example, right click **7uee_B** and click **set as reference**. Axes showing the directions of the first three principle components are drawn from this reference point. Click on the arrow or the very tip of the arrow (web version) pointing towards the other P1 branch and click on **explore axis**.

This will bring up a structure of your reference, **7uee_B**, and load the torsion angle changes associated with one of the principle component analysis (PCA) axes. If there are torsion angles missing from the **7uee_B** structure then a dialogue window will give an outline of the problem. It is good to manually check for disordered regions or deletions/insertions which may distort the motion in the middle of the protein backbone. Torsion angles missing from the N- or C-terminus, or in sidechains, are less detrimental to the overall motion of the protein. Press **OK** to dismiss the warning.

The slider at the bottom allows you to slide the position in RoPE space along your chosen PCA axis, ranging from −1 to 0 to +1. The colours of the torsion angle are proportional to the total change in Ramachandran angles for each Cα atom. Brightly coloured residues therefore denote hinge regions in the structure. Note that the torsion angle motion does not correspond to any single motion between two structures, but uses the information across all datasets to derive the principle independent motions. It is also displayed independently of any partially correlated motions which may accompany such a transition. The same RoPE axis will also produce different atomic motions depending on the reference structure. However, inspection of this motion shows a collapse of the active site of the protein. Reference to an external model-viewing program suggests a collapse of the distance between residues S118 and P296 across both sides of the active site to produce a closed conformation.

Let us examine this distance as a potential marker more carefully: the distance between S118 and P296 is a function of all backbone torsion angles in between these two residues. RoPE contains the ability to measure distances between corresponding atoms in two structures. Click **back** until returned to the **Entity - alcohol-dehydrogenase** menu and click **Sequence**. This loads the sequence from the UniProt ID. Use the left and right arrows in the screen to flick through successive pages. Click on residue 118 (**S**) and click **CA** (the Cα of residue 118), and click **Yes** to confirm you wish to measure from this atom. Flick to residue 296 residue CA (**P**) and confirm measurement to this atom. Wait for the distances to be calculated and then click **Add to database**.

You can now return to the RoPE space. Go to the **rules** menu, and **(+) add rule** to vary colour on the **118CA to 296CA** distance. **Create** this and return to the RoPE space. It is immediately visible that there is wide variation across all structures of this reporter distance. Structures which are not in P1 are more likely to have wider distances, with some exceptions. Firstly, there is far more variation across the lengths of the V branches than between them. The tips of the P1-V contain the lowest distances of about 12.2 Å. However, comparison of monomers from each model show a systematic increase in this reporter distance between chains assigned A to chains assigned B. This therefore demonstrates that when the protein crystallises as a closed-form dimer, there are nevertheless subtle and consistent differences between the subunits in this space group.

Additional walkthroughs are available on the website, including how to import locally saved PDB files, and introducing custom metadata.

### 6 Pitfalls for data interpretation

The number of PCA axes vastly exceeds the number displayable at any one time. The first three most important axes are displayed by default, but the researcher should always bear in mind additional separation between structures on non-displayed axes. The displayed axes can be altered in the **axes** menu when viewing the RoPE space.

In this version of RoPE, disordered regions in proteins are a source of breaking the correlation in the backbone. Ordered regions of the backbone are separately superimposed during PCA axis exploration as a reasonable alternative, but must be taken into account when evaluating motions.

How many protein structures are needed for the analysis? This depends on the scientific question one is trying to answer and therefore how much sensitivity is needed. The success of disentanglement of independently varying motions, particularly separation of the salient and ambient motions, relies on good sampling and coverage of the conformational space of the protein. Repeating experiments under identical conditions is key to understanding the natural variation of the sample. Make use of the group selection feature to eliminate modes of motion that are irrelevant to the biological question, which may otherwise pollute the PCA result.

### 7 Conclusions

The software allows the researcher to easily access the analysis described in this paper for their own projects and proteins, and provides a framework in which additional analytical tools will be made available for biological interpretation. Additional features not described in this paper are documented on the RoPE website.

## 8 Acknowledgements

Thank you to Dr David Briggs, Dr Gianluca Santoni, Professor David Stuart, Dr Diana Monteiro, Dr Benjamin Krishna and Dr Helen Duyvesteyn for a critical reading through the manuscript, and to Dimitris Triandafillidis for proofreading, helpful discussions, test cases and bug reports. The author reports no conflicts of interest.

## 9 Methods

### Definition and population of a protein entity

The UniProt sequence was obtained for trypsin and lysozyme, and the sequences downloaded and referred to as an entity. Sequence similarity was used to search for and download similar protein structures from the PDB. Alternatively, PDB codes associated with a single multi-dataset research project were downloaded for BAZ2B [12], PDB group deposition code G_1002036 and the time-resolved myoglobin CO-dissociation experiment [21] from PDB-redo [19]. The downloaded sequences from the PDB file were aligned against the entity reference sequence, identifying point mutations and correcting the register for insertions and deletions where necessary. Chains identified as having at least 10 residues and 80% identity to a target entity sequence were assigned as a member of the entity group. The mapping of residues to the reference sequence was recorded for each participating chain.

### Refinement and matching of torsion angles

Bond lengths and angles were reset to the best estimates from the CCP4 monomer library [32]. The initial torsion angles estimated from the atomic coordinate structure were then optimised using the first step of the Vagabond algorithm as previously described [20] to best match the model atomic coordinates. Torsion angles were defined by the four atom names which form the relationship (e.g. N-CA-C-N) as defined in the PDB file. The torsion angle was assigned to the residue of the middle two atoms. If the two atoms spanned two residues (e.g. a peptide bond or disulphide bond), the torsion angle was assigned to the earlier residue in the sequence. Torsion angles between two molecules were considered to have the same identity if the four atoms correspond in either their forward or reverse direction (N-CA-C-N or N-C-CA-N) and they were assigned to the same reference sequence residue. A per-molecule vector was constructed, comprising all unique torsion angle identities, per residue, across all molecules. If a molecule did not have a corresponding torsion angle, the value was set to not a number (nan) in memory. For example, torsion angles between two molecules differing by a point mutation will only be compared as far as their atom names remain congruent (e.g. lysine and arginine torsion angles will be comparable up to the delta carbon).

### Clustering of torsion angles

Torsion angle differences were calculated with respect to the average for each identified torsion angle across all participating datasets. Each torsion angle difference *t* was converted into a two-dimensional vector (*sin*(*t*), *sin*(*t*)*cos*(*t*)). Unassigned angles acquired a unit vector of of length zero. The cluster4x algorithm is applied to the differences in this two-dimensional vector format. A correlation matrix was generated by calculating correlation coefficients between matched torsion angles between each pair of data sets. Singular value decomposition (SVD), as implemented in Numerical Recipes in C [33], was applied to the resulting matrix, and the three top axes of variation were plotted at rotation angles which clearly show the variation in the datasets. Each point was remapped into the subspace generated using SVD and scaled on each axis according to the respective eigenvalues (equivalent to the variance of data when projected onto each axis). The graphical user interface allowed structures to be coloured by metadata categories with numerical values such as collection temperature or resolution. Marker types for structures were changed according to discrete metadata such as space group or point mutation, and series of related datasets were connected by lines. The full workflow for handling torsion angles is shown in Figure 2.

### Clustering of atomic coordinates

Clustering was also carried out using atomic coordinate differences in order to compare with the results obtained using torsion angles. In this case, usually, a reference molecule was chosen that had the highest number of modelled residues without exceeding the sequence length. However, in some situations, this automatically chosen reference was not suitable and an alternative was manually chosen. All other molecules were superposed onto the reference molecule using the Kabsch algorithm [34]. The Atomic coordinates were extracted for each atom, matched by atom name and sequence alignment. Clustering on the differences of atomic coordinates was carried out using the cluster4x algorithm and displayed using the same graphical user interface as for torsion angles.

## Notes

https://rope.hginn.co.uk

https://www.github.com/helenginn/rope

### Competing Interest Statement

The authors have declared no competing interest.

### Summary of Updates

Revised in light of referee comments, with updates to the methodology and comparison with atomic coordinates.

